# Discrete Roles of the Ir76b Ionotropic Co-Receptor Impact Olfaction, Blood Feeding, and Mating in the Malaria Vector Mosquito *Anopheles coluzzii*

**DOI:** 10.1101/2021.07.05.451160

**Authors:** Zi Ye, Feng Liu, Huahua Sun, Adam Baker, Laurence J. Zwiebel

**Affiliations:** Department of Biological Sciences, Vanderbilt University, 465 21^st^ Avenue South, Nashville, TN 37235, USA; Department of Plant Pathology, College of Agriculture, Guizhou University, Guiyang, China

**Keywords:** *Anopheles coluzzii*, Olfaction, Ionotropic receptor, Electrophysiology, Blood feeding.

## Abstract

Anopheline mosquitoes rely on their highly sensitive chemosensory apparatus to detect diverse chemical stimuli that drive the host-seeking and blood-feeding behaviors required to vector pathogens for malaria and other diseases. This process incorporates a variety of chemosensory receptors and transduction pathways. We have used advanced *in vivo* gene-editing and -labelling approaches to localize and functionally characterize the ionotropic co-receptor *AcIr76b* in the malaria mosquito *Anopheles coluzzii*, where it impacts both olfactory and gustatory systems. *AcIr76b* has a broad expression pattern in female adult antennal grooved pegs, T1 and T2 sensilla on the labellum, stylets, and tarsi, as well as the larval sensory peg. *AcIr76b* is co-localized with the Orco odorant receptor (OR) co-receptor in a subset of cells across the female antennae and labella. In contrast to *Orco* and *Ir8a*, chemosensory co-receptors that appear essential for the activity of their respective sets of chemosensory neurons in mosquitoes, *AcIr76b^-/-^* mutants maintain wild-type peripheral responses to volatile amines on the adult palps, labellum, and the larval sensory cone. Interestingly, *AcIr76b^-/-^* mutants display significantly increased responses to amines in antennal grooved peg sensilla while coeloconic sensilla reveal significant deficits in responses to several acids and amines. Behaviorally, *AcIr76b* mutants manifest significantly female-specific insemination deficits and, although *AcIr76b^-/-^* mutant females are able to locate, alight, and probe artificial blood hosts, they are incapable of blood feeding successfully. Taken together, our study reveals a multi-dimensional functionality of Ir76b in Anopheline olfactory and gustatory pathways that directly impacts the vectorial capacity of these mosquitoes.

**Summary:** Chemosensory receptors play crucial roles across mosquito lifecycles where they often form functional complexes that require cognate co-receptors. To better understand mosquito chemosensory pathways in the malaria vector mosquito *An. coluzzii* we have utilized advanced gene editing approaches to localize and functionally characterize the ionotropic receptor co-receptor *AcIr76b*. Expression of *AcIr76b* was observed in antennal grooved pegs and other accessory olfactory appendages. Mutagenesis of *AcIr76b* uncovers both reduced and elevated neuronal responses to amines, which suggests a role in response modulation. In addition to olfactory phenotypes, *AcIr76b* mutants display significantly impaired mating and blood feeding capabilities. Our data reveals discrete roles of *AcIr76b* across olfactory and gustatory pathways and shed lights on the potential molecular target for vector control strategies.

## Introduction

The malaria mosquito *Anopheles coluzzii* (recently renamed from the “M” form of *An. gambiae* (1)) is a major vector of human malaria pathogens in subSaharan Africa that are transmitted during blood feeding by adult females (2). Anophelines and other mosquitoes locate blood-meal hosts through detection of a variety of environmental and host-derived cues, among which olfactory signals have great significance at both long and short range (3, 4). On the head of adult mosquitoes, the primary olfactory appendages include the antennae, maxillary palps and labella, which are covered by a range of hair-like protrusions, known as sensilla (5–8). One or more bipolar olfactory sensory neurons (OSNs) innervate a typical chemosensory sensillum where their dendrites extend apically and are bathed within an aqueous lymph (8, 9). Three large gene families encode the distinct molecular receptors that underlie olfaction in mosquitoes and other insects; these include odorant receptors (ORs), ionotropic receptors (IRs), and gustatory receptors (GRs) (4). These receptors are expressed on the dendritic membranes of diverse sets of chemosensory neurons, where they generate action potentials in response to a broad spectrum of chemical stimuli (5, 7, 10–13). Three major morphological types of olfactory sensilla are present on the mosquito antennae: trichoid, basiconic (also known as grooved peg), and coeloconic sensilla (9, 14). In *An. coluzzii*, while there is a range of functional variations within each type of sensillum, trichoid sensilla generally respond to a broad spectrum of odorants, while both grooved pegs and coeloconic sensilla appear to be more narrowly tuned to both amines and acids (6, 15, 16).

While molecularly unrelated, mosquito ORs and IRs are both ligand-gated heteromeric channels composed of tuning subunits along with one or more highly conserved co-receptor subunits (17–20). The tuning subunit is responsible for the specificity of the receptor while the co-receptor maintains the structural integrality and is crucial for the receptor function (19–21). Knockout (null) mutants of the odorant receptor co-receptor (Orco) in *Aedes aegypti* and *An. coluzzii* result in a dramatic decrease in sensitivity to a variety of human and other odorants; however, importantly, humans remain attractive to host-seeking *Orco^-/-^* mutants, highlighting the involvement of other odorant signaling pathways and sensory modalities (21, 22).

In *Drosophila*, antennal IRs are primarily expressed in the coeloconic sensilla (18). Unlike ORs, which rely solely on the Orco co-receptor, three IRs—Ir8a, Ir25a, and Ir76b—function as IR co-receptors (18, 19). Interestingly, each DmIR co-receptor is associated with distinct odor preferences; *DmIr8a* is critical for acid sensitivity, whereas *DmIr25a* and *DmIr76b* are both responsible for amine detection (18, 23, 24). In addition to olfactory function, Ir76b and Ir25a are also involved in other sensory modalities and pathways. In *Drosophila*, DmIr76b has been found to be involved in gustatory responses to salt and amino acids (25, 26), and DmIr25a acts in both thermosensation and hydrosensation (27, 28). Furthermore, while *DmIr25a* and *DmIr76b* are both co-expressed with DmIr92a acting as a *Drosophila* ammonia/amine receptor, DmIr92a is able to function independently of either these co-receptors (24, 29).

Recently, several studies focusing on mosquito IRs have revealed that the homologs to *Drosophila* IR co-receptors similarly regulate a range of Orco-independent sensing pathways. In the arbovirus vector *Ae. aegypti*, *AeIr8a* null mutants lost the neuronal and behavioral responses to acids (30). In *An. coluzzii* larvae, all three IR co-receptors are expressed on the larval antennae, and RNAi knockdown of *AcIr76b* specifically impacts larval responses to butylamine (31). In adults, *AcIr76b* is highly expressed in antennal neurons that do not express ORs (11); in *Xenopus* oocytes it drives responses to several amines when co-expressed with *AcIr25a* and either *AcIr41a* and *AcIr41c* (11).

To examine the roles and relevance of Ir76b in the olfactory system of *An. coluzzii*, we have utilized CRISPR/Cas9-mediated gene editing to establish an *AcIr76b*-*QF2* driver and *AcIr76b* null mutant lines. Localization studies using the driver within a binary Q system (32) reveal that *AcIr76b* is robustly expressed in antennal grooved pegs, labella, stylets and tarsi of adult females, as well in the larval antennae where it specifically innervates the sensory peg. Surprisingly, adult female *AcIr76b^-/-^* mosquitoes display significantly increased antennal responses to several amines, whereas peripheral responses to acid stimuli are unaffected. Behaviorally, *AcIr76b* mutant females display severe mating deficits and, interestingly, have acutely lost the ability to blood feed successfully. These studies demonstrate that *AcIr76b* acts in both olfactory and gustatory systems of *An. coluzzii* where it impacts the reproductive fitness and ultimately the vectorial capacity of this globally important mosquito.

## Materials and Methods

### Mosquito rearing

*An. coluzzii* (SUA 2La/2La), previously known as *Anopheles* gambiae sensu stricto “M-form” (1), originating from Suakoko, Liberia, were reared using previously described protocols. Briefly, all mosquito lines were reared at 27°C, 75% humidity under a 12h light/12h dark photoperiod and supplied with 10% sucrose water in the Vanderbilt University Insectary (33, 34). For stock propagation, 5- to 7-day-old mated females were blood fed for 30-45 min using a membrane feeding system (Hemotek, Lancaster, UK) filled with defibrinated sheep blood purchased from Hemostat Laboratories (Dixon, CA). Mosquito larvae were reared in distilled water at 27°C under the standard 12h light/12h dark cycle, with approximately 300 larvae per rearing pan in 1L H_2_O. The larval food was made from 0.12g/mL Kaytee Koi’s Choice premium fish food (Chilton, WI) plus 0.06g/mL yeast in distilled water and subsequently incubated at 4°C overnight for fermentation. For first and second instar larvae, 0.08mL larval food was added into the water every 24h. The *An. coluzzii* effector line (QUAS-mCD8:GFP) was a generous gift from the lab of Dr. C. Potter at The Johns Hopkins University School of Medicine.

### Mosquito mutagenesis

CRISPR/Cas9 gene editing in *An. coluzzii* was carried out as previously described (35). The CRISPR gene-targeting vector was a kind gift from the lab of Dr. Crisanti at Imperial College London (36). The single guide RNA (sgRNA) sequence was designed by CHOPCHOP (37) to target the first exon of *AcIr76b* gene (ACOM032257). The complimentary oligos (Ir76b_gRNA_F/Ir76b_gRNA_R; **Table S1**) were artificially synthesized (Integrated DNA Technologies, Coralville, IA) and subcloned into the CRISPR vector by Golden Gate cloning (New England Biolabs, Ipswich, MA). The homologous templates were constructed based on a pHD-DsRed vector (a gift from Kate O’Connor-Giles; Addgene plasmid #51434; http://n2t.net/addgene:51434; RRID:Addgene 51434) where the 2-kb homologous arms extending either direction from the double-stranded break (DSB) site were PCR amplified with the customized oligonucleotide primers: Ir76b_AarI_F/Ir76b_AarI_R; Ir76b_SapI_F/Ir76b_SapI_R (**Table S1**) and sequentially inserted into the AarI/SapI restriction sites on the vector.

The microinjection protocol follows a previous study (16, 38). In brief, newly laid (approximately 1h-old) embryos of the wild-type *An. coluzzii* were immediately collected and aligned on a filter paper moistened with 25mM sodium chloride solution. All the embryos were fixed on a coverslip with double-sided tape, and a drop of halocarbon oil 27 was applied to cover the embryos. The coverslip was further fixed on a slide under a Zeiss Axiovert 35 microscope with a 40× objective. The microinjection was performed using Eppendorf FemtoJet 5247 (Eppendorf, Hamburg, Germany) and quartz needles (Sutter Instrument, Novato, CA). The gene-targeting vector and the homologous template were co-injected at 300ng/μL each. Injected embryos were subsequently placed in deionized water with artificial sea salt (0.3g/L) and thereafter reared under normal VU insectary conditions.

First generation (G0) of injected adults were separated based on gender and crossed to 5× wild-type gender counterparts. Their offspring (F1) were screened for DsRed-derived red eye fluorescence. Red-eyed F1 males were individually crossed to 5× wild-type females to establish a stable mutant line. DNA extraction was performed using DNeasy Blood & Tissue kits following the manufacturer’s instruction (Qiagen, Hilden, Germany) and the genomic DNA was used as templates for PCR analyses of all individuals (after mating) to validate the fluorescence marker insertion using primers that cover double-stranded break site (Ir76b_F/Ir76b_R; **Table S1**). Salient PCR products were sequenced to confirm the accuracy of the genomic insertion. The heterozygous mutant lines were thereafter back-crossed to the wild-type *An. coluzzii* for at least five generations before putative homozygous individuals were manually screened for DsRed-derived red eye fluorescence intensity. Putative homozygotic mutant individuals were mated to each other before being sacrificed for genomic DNA extraction and PCR analyses (as above) to confirm their homozygosity.

To generate the *AcIr76b-QF2* driver line for Q system, *T2A-QF2-3xP3-DsRed* element was inserted into the *Ir76b* coding region through CRISPR-mediated homologous recombination. DSB was induced using the CRISPR gene-targeting vector described above. The homologous template that contains *T2A-QF2-3xP3-DsRed* element, which was amplified from the ppk301-T2A-QF2 HDR plasmid (Matthews et al., 2019; A gift from Leslie Vosshall; Addgene plasmid# 130667; http://n2t.net/addgene:130667; RRID:Addgene_130667), flanked by ∼2kb homologous arms, was constructed using NEBuilder HiFi DNA Assembly kit (NEB, Ipswich, MA) (Ir76b_LArm_F/Ir76b_LArm_R; Ir76b_RArm_F/Ir76b_RArm_R; Ir76b_T2A_F/Ir76b_T2A_R; **Table S1**). From the left homologous arm immediately preceding the *T2A*, 2bp was removed to keep the *T2A* sequence in-frame. Red-eyed F1 mosquitoes were backcrossed for three generations and then crossed to the effector line to acquire progeny for Ir76b localization studies.

### Immunohistochemistry

Antibody staining was performed as previously described (10, 16). Antennae, labella, and maxillary palps were dissected into 4% formaldehyde in PBST and fixed on ice for 30min. Samples were washed 3× in PBST for 10min each and then embedded in TFM (Tissue Freezing Medium; General Data Company Inc., Cincinnati, OH). Cryosections were obtained at −20°C using a CM1900 cryostat (Leica Microsystems, Bannockburn, IL). Samples were sectioned at ∼10μm and transferred onto Superfrost plus slides (VWR Scientific, Radnor, PA). Slides were air-dried at room temperature (RT) for 30min and fixed in 4% formaldehyde in PBST for 10min, followed by 3× rinsing in PBST for 10min each. Thereafter, 5% normal goat serum (Sigma-Aldrich, St. Louis, MO) in PBST was applied and the slides were blocked in the dark at RT for 1h in HybriWell sealing chambers (Grace Bio-Labs, Bend, OR). Primary antibody Rabbit α-Orco was diluted 1:500 in 5% normal goat serum in PBST and applied on the slides and incubated overnight at 4°C.

After primary antibody staining, slides were washed 3× in PBST for 10min each and stained with secondary antibody Goat α-Rabbit-Cy3 (Jackson ImmunoResearch, West Grove, PA) 1:500 in 5% normal goat serum PBST for 2h at RT and then rinsed 3×. Nuclei were stained with 300nM DAPI (Invitrogen, Carlsbad, CA) at RT for 10min. Slides were briefly washed and mounted in Vectashield medium (Vector Laboratories, Burlingame, CA). Whole-mount samples were dissected into 4% formaldehyde in PBST and fixed on ice for 30min. After 3× washing in PBST for 10min each, the samples were transferred onto slides and mounted in Vectashield medium (Vector Laboratories, Burlingame, CA). Confocal microscopy images at 1024×1024 pixel resolution were collected from under an Olympus FV-1000 equipped with a 100× oil objective at the Vanderbilt University Cell Imaging Shared Resource Core. Laser wavelengths of 405nm, 488nm, and 543nm were used to detect DAPI, GFP, and Cy3, respectively.

### Transcuticular electrophysiology

Electroantennogram (EAG), electropalpogram (EPG), and electrolabellogram (ELG) recordings were conducted following a previous study, with modifications (7, 34). Female mosquitoes (4-10 days after eclosion) with legs and wings removed were fixed on glass slides using double-sided tape. The last segment of antenna with the tip partially cut was subsequently connected to a recording glass electrode filled with Ringer solution (96mM NaCl, 2mM KCl, 1mM MgCl_2_, 1mM CaCl_2_, 5mM HEPES, pH = 7.5), to which a tungsten wire was in contact to complete a circuit with another glass reference electrode similarly connected into the compound eye of the female. The antennal preparation was continuously exposed to humidified, charcoal-filtered air flow (1.84 L/min) transferred through a borosilicate glass tube (inner diameter = 0.8cm) using a stimulus controller (Syntech, Hilversum, The Netherlands), and the open end of the glass tube was located 1 cm from the antennal preparation. All chemicals were diluted to 10^−1^ (v/v) working solutions in paraffin oil except for lactic acid, acetic acid, ammonia, and dimethylamine which were diluted in ddH_2_O. 10-μl aliquots of test or control stimuli were transferred onto a piece of filter paper (3 × 50 mm) which was then placed inside the Pasteur pipette. Odor was delivered to the antennal preparation for 500ms through a hole placed on the side of the glass tube located 10cm from the open end of the tube (1.08 L/min), and the stimulus odor was mixed with continuous air flow through the hole. A charcoal-filtered air flow (0.76 L/min) was delivered from another valve through a blank pipette into the glass tube at the same distance from the preparation in order to minimize changes in flow rate during odor stimulation. The resulting signals were amplified 10× and imported into a PC via an intelligent data acquisition controller (IDAC, Syntech, Hilversum, The Netherlands) interface box, and then recordings were analyzed using EAG software (EAG Version 2.7, Syntech, Hilversum, The Netherlands). Response amplitudes were normalized by dividing the response amplitude of odorant stimuli by the response amplitude of control (solvent alone) responses.

### Single sensillum recording

Single sensillum recordings (SSR) were carried out as previously described (40, 41) with minor modifications. Female adult mosquitoes (4-10 days after eclosion) were mounted on a microscope slide (76 × 26 mm). Using double-sided tape, the antennae were fixed to a cover slip resting on a small bead of dental wax to facilitate manipulation, and the cover slip was placed at approximately 30 degrees to the mosquito head. Once mounted, the specimen was placed under an Olympus BX51WI microscope and the antennae viewed at high magnification (1000×). Recordings were carried out using two tungsten microelectrodes freshly sharpened in 10% KNO_2_ at 10V. The grounded reference electrode was inserted into the compound eye of the mosquito using a WPI micromanipulator, and the recording electrode was connected to the preamplifier (Syntech universal AC/DC 10×, Syntech, Hilversum, The Netherlands) and inserted into the shaft of the olfactory sensillum to complete the electrical circuit to extracellularly record OSN potentials (42). Controlled manipulation of the recording electrode was performed using a Burleigh micromanipulator (Model PCS6000). The preamplifier was connected to an analog-to-digital signal converter (IDAC-4, Syntech, Hilversum, The Netherlands), which in turn was connected to a computer for signal recording and visualization.

All chemicals were diluted to 10^−2^ (v/v) working solutions in paraffin oil except for lactic acid, acetic acid, and dimethylamine which were diluted in ddH_2_O. For each chemical, a 10-μl aliquot was applied onto a filter paper (3 × 50mm) which was then inserted into a Pasteur pipette to create the stimulus cartridge. A sample containing the solvent (paraffin oil or water) alone served as the control. The airflow was maintained at a constant 20mL/s throughout the experiment. Purified and humidified air was delivered to the preparation through a glass tube (10-mm inner diameter) perforated by a small hole 10cm away from the end into which the tip of the Pasteur pipette could be inserted. The stimulus was delivered to the sensilla by inserting the tip of the stimulus cartridge into this hole and diverting a portion of the air stream (0.5L/min) to flow through the stimulus cartridge for 500ms using a stimulus controller (Syntech, Hilversum, The Netherlands). The distance between the end of the glass tube and the antennae was ⩽1cm. Signals were recorded for 10s, starting 1s before stimulation, and the action potential spikes were counted off-line over a 500-ms period before and after stimulation. Spike rates observed during the 500-ms stimulation were subtracted from the spontaneous (background) spike activity observed in the preceding 500ms, and counts were recorded in units of spikes/s. Post-stimulus “OFF” responses were calculated by subtracting the pre-stimulus background activities from the spike counts in the 1s immediately following the stimulus application. Responses of odorants were always normalized by subtracting the solvent responses.

### Mating bioassay

Newly emerged wild-type females and males were separated from day 1 to ensure no mating occurred. 15 females and 10 males were then placed in a rearing bucket and allowed to mate freely for 5 days. All surviving females were then collected and their spermathecae were dissected under a compound microscope. The spermathecae were then placed in the buffer (145mM NaCl, 4mM KCl, 1mM MgCl2, 1.3mM CaCl2, 5mM D-glucose, 10mM HEPES) (43) with 300nM DAPI, and a cover slip was used to gently press and break the spermathecae to release the sperm. As in previous studies (15), the spermathecae were examined, under the 1000× compound microscope, to assess the insemination status. The insemination rate was calculated by dividing the number of inseminated females by the total number of females in each bucket.

### Blood feeding bioassay

Feeding bioassays were carried out for the 6- to 8-day-old wild-type and *AcIr76b* females between ZT11 and ZT12 during which *An. coluzzii* was shown to be behaviorally active. Mosquitoes were starved, with only water access, for 24h prior to the bioassay. Defibrinated sheep blood was stored at 4°C and used within 2 weeks of purchasing (Hemostat Laboratories, Dixon, CA). Human foot odorants were provided as cloth strips cut from socks that had been continually worn for 5 days by a 30-year-old male volunteer and thereafter incubated overnight at 37°C in a sealed Ziploc plastic bag (SC Johnson, Racine, WI). For each replicate, 25-30 female mosquitoes were released into a 32-oz container, the top of which was covered with a net through which the mosquito proboscises are able to penetrate to feed but not to escape. A membrane feeder (Hemotek, Lancaster, UK) that heated the blood meal to 37°C was then placed on the net. A mini camera (GoPro, San Mateo, CA) was set at the bottom of the container to record the landing activity of mosquitoes. A human volunteer smoothly exhaled into the container for 5 s to activate the mosquitoes with CO_2_ for blood feeding. The mosquitoes were allowed to feed freely for 25min and then were immediately anesthetized at −20°C in order to assess the number with blood-engorged abdomens, indicative of successful blood feeding. The assay containers and videos were also analyzed post hoc by volunteers blinded to the experimental genotypes, who manually counted both blood-engorged mosquitoes and the number of landings onto the feeder during the assay. The total landing count was divided by the total mosquito number in each assay to calculate the landing number per mosquito.

### Capillary feeder (CAFE) bioassay

The CAFE bioassay was conducted following a previous study, with minor modifications (15, 44). Each trial started at ZT12 and ended at ZT18 for 6h. Four 4- to 8-day-old mosquitoes were provided with water but otherwise fasted for 22h before being anesthetized on ice briefly and placed into a *Drosophila* vial (24.5 × 95mm; Fisher Scientific, Waltham, MA). A borosilicate glass capillary (1B100F-3; World Precision Instruments, Sarasota, FL) was filled with 10% sucrose water and embedded into a cotton plug. The vial opening was then blocked with the cotton plug and the capillary was placed slightly protruding from the plug into the vial for mosquitoes to feed on. The sugar level in the capillary was compared before and after each trial to generate the initial sugar consumption value. At least four control vials with no mosquitoes inside were used to assess the evaporation at the same time. The final sugar consumption was calculated by subtracting the evaporation from the initial sugar consumption value.

## Results

### *AcIr76b* expression in antennal grooved pegs

In order to identify the types of antennal sensilla in which AcIr76b activity is salient, the spatial expression pattern of *AcIr76b* was established using the binary expression Q system (16, 45), which provides more sensitivity than fluorescence *in-situ* hybridization (FISH)-based methods previously used (11). This system requires genetic crosses between a QUAS-GFP effector line and a *AcIr76b promoter*-*QF2* (*AcIr76b-QF2*) driver line, resulting in GFP fluorescence in *AcIr76b*-expressing cells. While previous Anopheline driver lines were developed by integrating *promoter-QF2* constructs into pre-defined or random genomic sites (16, 45), those promoters were presumptive as they were derived from selected regions of 5’ upstream sequences without precise elucidation of the required regulatory sequences. As such, those promoters are potentially error-prone and artifactual, leading to localizations that are compromised by over-/under-expression from fragmented or otherwise partial regulatory information. To overcome this limitation, we incorporated a newly developed approach using CRISPR-mediated homologous recombination to insert a *T2A-QF2-3xP3-DsRed* element into the *AcIr76b* locus at the first exon (**Figure 1A**) (39), and the knock-in was subsequently confirmed by means of genomic PCR (**Figure 1B**) and sequencing. The T2A peptide induces ribosomal skipping which facilitates the unbiased expression of *QF2* driven by endogenous, fully intact *AcIr76b* regulatory sequences (46). For localization studies, F1 progeny derived from crosses between appropriate parental driver and effector lines would therefore express the *QF2* transcriptional factor that specifically binds to the QUAS activation sequence to drive expressions of the visual marker *GFP* in all *An. coluzzii* cells that normally express *AcIr76b*.

**Figure 1.**
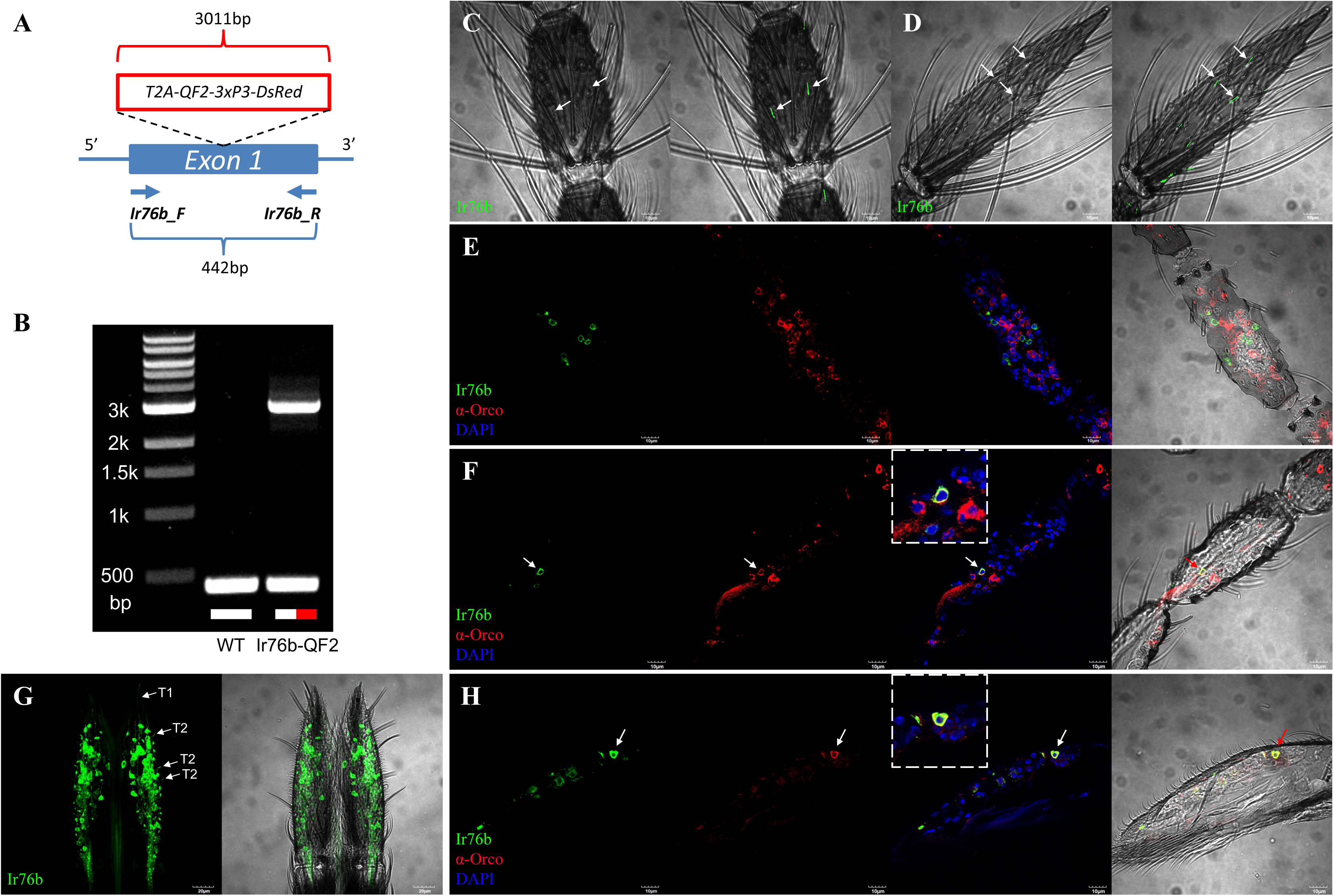
**(A)** *AcIr76b-QF2* driver line schematics. The *T2A-QF2-3xP3-DsRed* element was inserted into the first exon of *AcIr76b* via CRISPR-mediated homologous recombination. A pair of primers (*Ir76b_F* and *Ir76b_R*) were used for genomic PCR validation. The wild type produces a 442-bp amplicon whereas the *AcIr76b-QF2* driver allele gives rise to a 3011-bp amplicon. **(B)** Agarose electrophoresis of genomic PCR validation of wild type (WT) and driver (Ir76b-QF2). **(C)** Representative bright-field image (left) with GFP fluorescence overlay confocal z-stack projects (right) of whole-mount female antennae indicating *AcIr76b* is expressed in grooved pegs on 6^th^ flagellomere and **(D)** 13^th^ flagellomere (highlighted by arrows). **(E)** A representative confocal optical section of the female antennae immunohistochemically labelled with **α-**Orco antisera (red) and *AcIr76b* (green) indicating that AcIr76b and Orco are localized in distinct cells. **(F)** A representative confocal optical section of a female antenna showing AcIr76b and Orco are co-localized in a cell (highlighted by an arrow and enlarged within dashed lines). **(G)** A representative confocal Z-stack project of a whole-mount female labellum showing *AcIr76b* is expressed in dendrites in T1 and T2 sensilla (highlighted by arrows. Scale bars = 20μm). **(H)** A representative confocal optical section of a female labellum showing AcIr76b and Orco are co-localized in a cell (highlighted by an arrow and enlarged within dashed lines). Nuclei were labelled with DAPI. Scale bars = 10μm.

In these studies, extensive *GFP* labelling was observed in antennal grooved pegs (basiconic sensilla) of 4- to 6-day-old adult female *An. coluzzii*, which would be expected to be actively seeking blood meals. Across multiple replicates that were examined on the 2^nd^-13^th^ antennal flagellomeres, 100% of grooved pegs, easily identified through their distinctive morphology, contained *AcIr76b*-expressing neurons displaying GFP-derived fluorescence in their dendrites (**Figure 1C&D**). In contrast, GFP signal was never observed in any trichoid sensilla screened across more than 20 individual antennal preparations. Recent studies have uncovered non-canonical co-expression of IRs and ORs in a subset of chemosensory neurons of *Drosophila* and *Ae. aegypti* (47, 48). In that light, we investigated whether IR/OR co-expression also occurs in *An. coluzzii* by immunolocalization using Orco-specific antibodies together with *Ir76b-QF* progeny. Our data suggest that while the majority (>95%) of *Ir76b*-*GFP*-positive cells are not labelled by Orco antibodies (**Figure 1E**), at least a small subset do indeed co-express *AcIr76b* and *Orco* (**Figure 1F**). Inasmuch as our examination of antennal trichoid sensilla as well as previous studies of *An. coluzzii* grooved pegs indicate they are devoid of Orco protein (10), the co-expression of *AcIr76b* and *Orco* seen here likely reflect either cryptic coeloconic or an as-yet unidentified population of sensilla.

In addition to the antennae of *An. coluzzii* females, *AcIr76b* localization analyses revealed that while the maxillary palps are devoid of *AcIr76b*-expressing cells, they are highly enriched in the labella (**Figure 1G**) and also present on the pre-, meso-, and meta-tarsi (**Figure S1**). On the *An. coluzzii* female labellum, *AcIr76b* is expressed in two populations of neurons with distinctive dendrites: the relatively long dendrites that innervate gustatory T1 sensilla and the short dendrites that are associated with olfactory T2 sensilla (**Figure 1G**) (7, 49). As was the case for the antennae, *AcIr76b* and *Orco* are co-expressed in a subset of cells present on the labella of *An. coluzzii* females (**Figure 1H**).

### Generation of *AcIr76b* null mutants

The *AcIr76b* null mutant line was generated using the CRISPR/Cas9 system together with the double-strand break (DSB)-specific homology template in order to knock-in a *3xP3-DsRed* visible marker construct at the *AcIr76b* DSB site. The sgRNA was chosen to target the first exon of *AcIr76b* to produce an early stop codon that leads to a malfunctional protein (**Figure 2A**). In total, 376 pre-blastoderm embryos were microinjected with the homologous template and the CRISPR targeting vector, in which *Cas9* expression is regulated by the *vasa2* promoter to specifically induce germ cell mutagenesis. Of the 17 larvae (4.5%) that survived the injection, 2 female and 2 male adults were able to successfully eclose and were designated as G0. These adults were then collectively crossed with the wild-type population, and the progeny (F1) were screened for the presence of red fluorescence in the eye and ventral nerve cord that would be expected if they contain a *3xP3-DsRed* insert. At least one of the G0-injected adults produced red-eyed F1 offspring, which indicates a high mutagenesis efficiency (≥25%). Red-eyed F1 males were individually back-crossed with wild-type females after which their *AcIr76b* mutant genotype was confirmed by genomic PCR (**Figure 2B**) and DNA sequencing. Homozygous mutants were generated by crossing heterozygotes with progeny selected based on their high intensities of red fluorescence, and thereafter genomic PCR (**Figure 2B**) and DNA sequencing was carried out to confirm phenotypic assessments.

**Figure 2.**
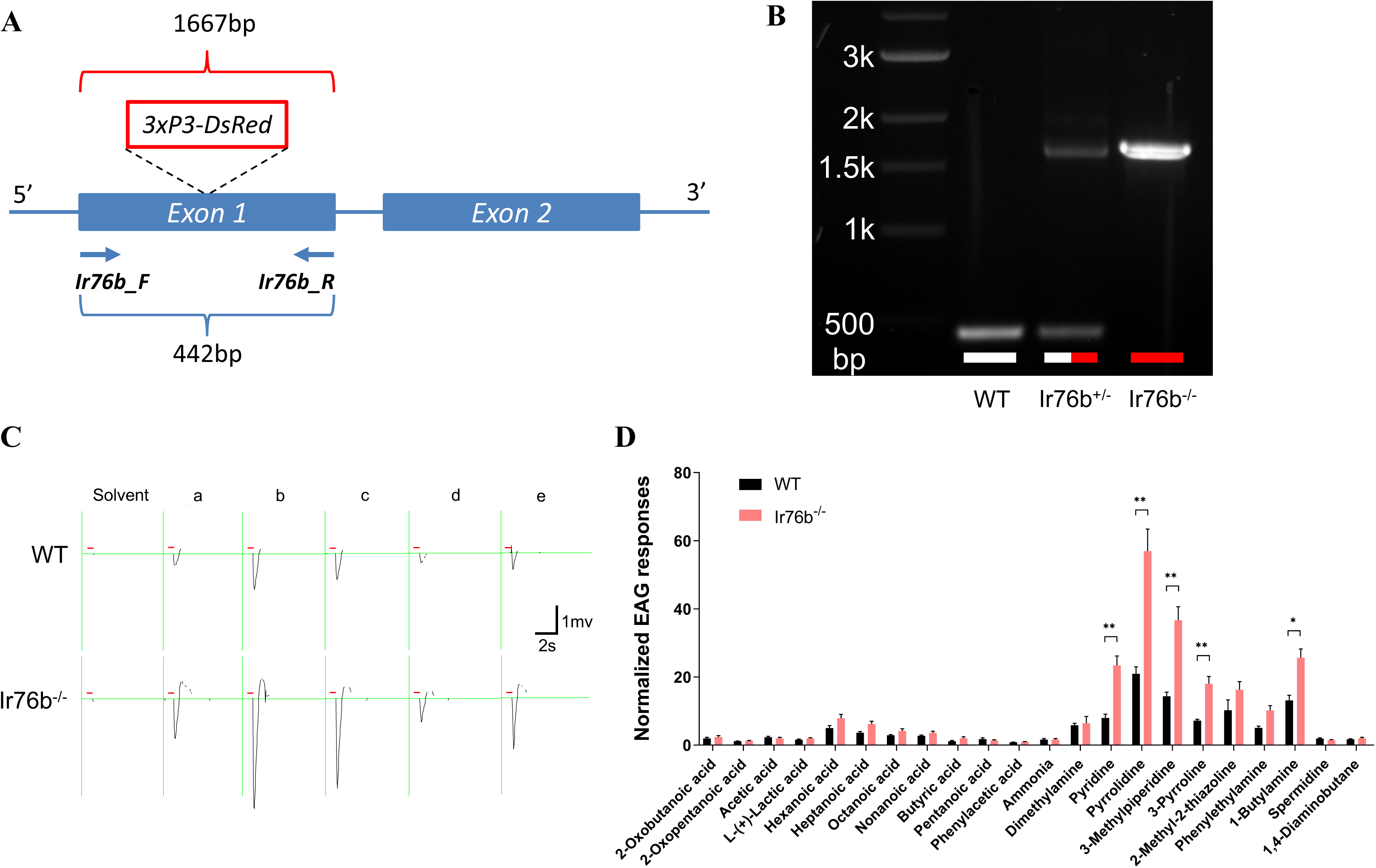
**(A)** *Ac*Ir76b mutagenesis schematics. The *3xP3-DeRed* element was inserted into the first exon of *AcIr76b* via CRISPR-mediated homologous recombination. A pair of primers (*Ir76b_F* and *Ir76b_R*) were used for PCR validation. The wild-type genome produces a 442-bp amplicon whereas the mutant allele gives rise to a 1667-bp amplicon. **(B)** Agarose electrophoresis of genomic PCR validation of wild type (WT), heterozygotes (Ir76b^+/-^), homozygotes (Ir76b^-/-^) templates. **(C)** Representative EAG recordings of wild-type (WT) and *AcIr76b^-/-^* (Ir76b^-/-^) females in response to paraffin oil (solvent) and a panel of amines including (a) pyridine, (b) pyrrolidine, (c) 3-methylpiperidine, (d) 3-pyrroline, and (e) 1-butylamine. The red bars indicate stimulus duration (0.5s). **(D)** Average EAG responses to a panel of acids and amines. Multiple t-tests using Holm-Sidak method (N=8) suggest responses to amines in *AcIr76b* mutants are significantly higher than in wild type. Responses were normalized to the solvent responses. Significance levels are depicted with asterisks: p-value < 0.05 (*); p-value < 0.01 (**); p-value < 0.001 (***). Error bars = Standard error of the mean.

### Electroantennogram revealed increased responses to amines in *AcIr76b* mutants

In order to initially investigate the function of AcIr76b in adult antennal olfactory responses, transcuticular EAG studies were carried out to broadly compare responses between *AcIr76b^-/-^* and wild-type females against a panel of acids and amines. Surprisingly, in contrast to response deficits typically seen in other olfactory co-receptor mutants studies (21, 22, 30), the EAG responses to several amines, including pyridine, pyrrolidine, 3-methylpiperidine, 3-pyrroline, and 1-butylamine, were significantly higher in *AcIr76b^-/-^* females than wild type (**Figure 2C&D**). In addition, EAG studies failed to reveal a significant impact on responses of *AcIr76b^-/-^* females to a panel of acid stimuli (**Figure 2D**).

### Elevated single sensillum responses in *AcIr76b^-/-^* grooved pegs

The localization of antennal *AcIr76b* suggests that the increased EAG responses to amines (**Figure 2C&D**) are most likely due to elevated neuronal responses in populations of grooved pegs. To test this hypothesis, SSR studies were carried out across a randomized sampling of 14 antennal grooved pegs to characterize and compare response profiles of *AcIr76b^-/-^* and wild-type females. The data reveal a range of wild-type responses to several amines that includes both excitation and inhibition (**Figure 3A&B**). Interestingly, while *AcIr76b^-/-^* mutants displayed wild-type levels of spontaneous (background) neuronal activity (**Figure 3C**) as well as responses to pyridine, 2-methyl-2-thiazoline, and phenylethylamine, we observed significantly elevated excitation elicited by pyrrolidine, 3-pyrroline, 3-methylpiperidine, and 1-butylamine compared with the wild-type responses (**Figure 3A, 3B & Figure S2**). Furthermore, post-stimulus “OFF” responses of grooved pegs were also analyzed, revealing significantly higher levels of neuronal spiking in the aftermath of 3-methylpiperidine stimulation in the mutants than in their wild-type counterparts (**Figure 3D**). No significant differences were detected in the olfactory responses to amines from antennal trichoid sensilla when comparing *AcIr76b^-/-^* mutants and similarly aged wild-type female *An. coluzzii* (**Figure 3E**).

**Figure 3.**
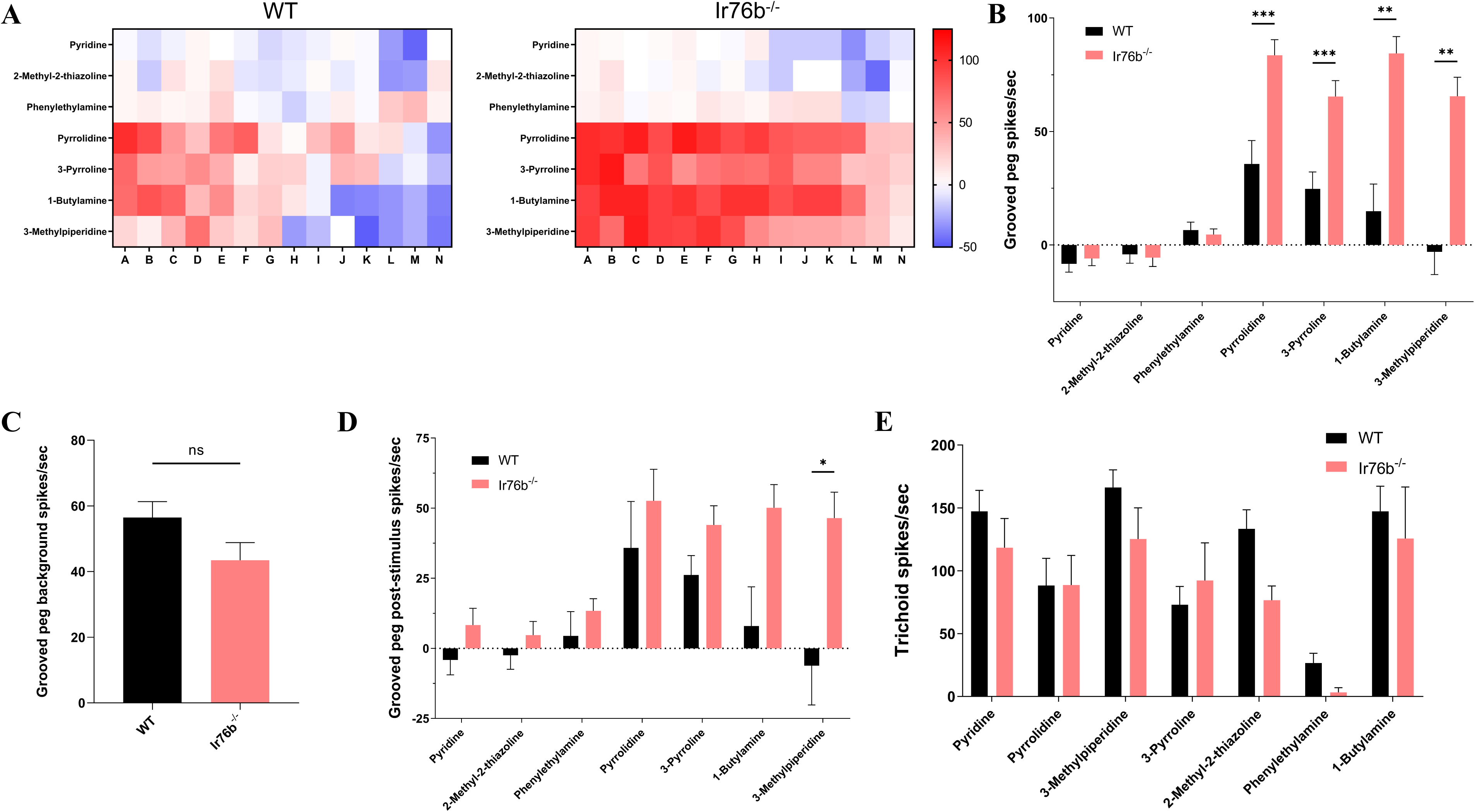
**(A)** Heatmaps showing average female grooved peg SSR responses to amines in the wild-type (WT) and *AcIr76b^-/-^* (Ir76b^-/-^) females. Each column depicts a single replicate. The color scale marks the response amplitude from the highest (125 spikes/s, red) to the lowest (−50 spikes/s, blue). Responses were normalized by subtracting the solvent responses. **(B)** Average female grooved peg SSR responses. Multiple t-tests using Holm-Sidak method (N=14) suggest responses to amines in *AcIr76b* mutants are significantly higher than in wild type. **(C)** Average female grooved peg spontaneous (background) neuronal activity. Non-parametric t-test suggests no significant differences between *AcIr76b* mutants and wild-type genotypes. **(D)** Average female grooved peg post-stimulus responses. Multiple t-tests using Holm-Sidak method (N=13) suggest post-stimulus “OFF” responses to 3-methylpiperidine in *AcIr76b* mutants are significantly higher than in wild type. **(E)** Average female trichoid SSR responses. Multiple t-tests using Holm-Sidak method (N=6) suggest no significant differences between *AcIr76b* mutants and the wild type. Significance levels are depicted with asterisks: p-value < 0.05 (*); p-value < 0.01 (**); p-value < 0.001 (***). Error bars = Standard error of the mean.

Considering that the technical challenges of visualization within the unique ultrastructure of peg-in-pit antennal coeloconic sensilla might account for the absence of GFP signals in *AcIr76b-GFP* whole-mounts, randomized SSR responses to acids and amines across the 2^nd^-8^th^ flagellomeres were also carried out to explore potential *AcIr76b* function (**Figure 4A**). While wild-type and *AcIr76b^-/-^* female coeloconic sensilla displayed similar levels of spontaneous (background) neuronal activity (**Figure 4B**), *AcIr76b^-/-^* females became indifferent to hexanoic acid, phenylacetic acid, pyridine, pyrrolidine, 2-methyl-2-thiazoline, and 1,4-diaminobutane in contrast to robust wild-type responses (**Figure 4C**). Similar *AcIr76b^-/-^* decreases in “OFF” responses were also observed after stimulation with phenylacetic acid and pyridine (**Figure 4D**). While *AcIr76b^-/-^* females exhibited modest, albeit non-significant increases in acetic acid responses compared with wild types (**Figure 4C**), these mutants displayed significantly higher post-stimulus “OFF” responses to acetic acid (**Figure 4D**).

**Figure 4.**
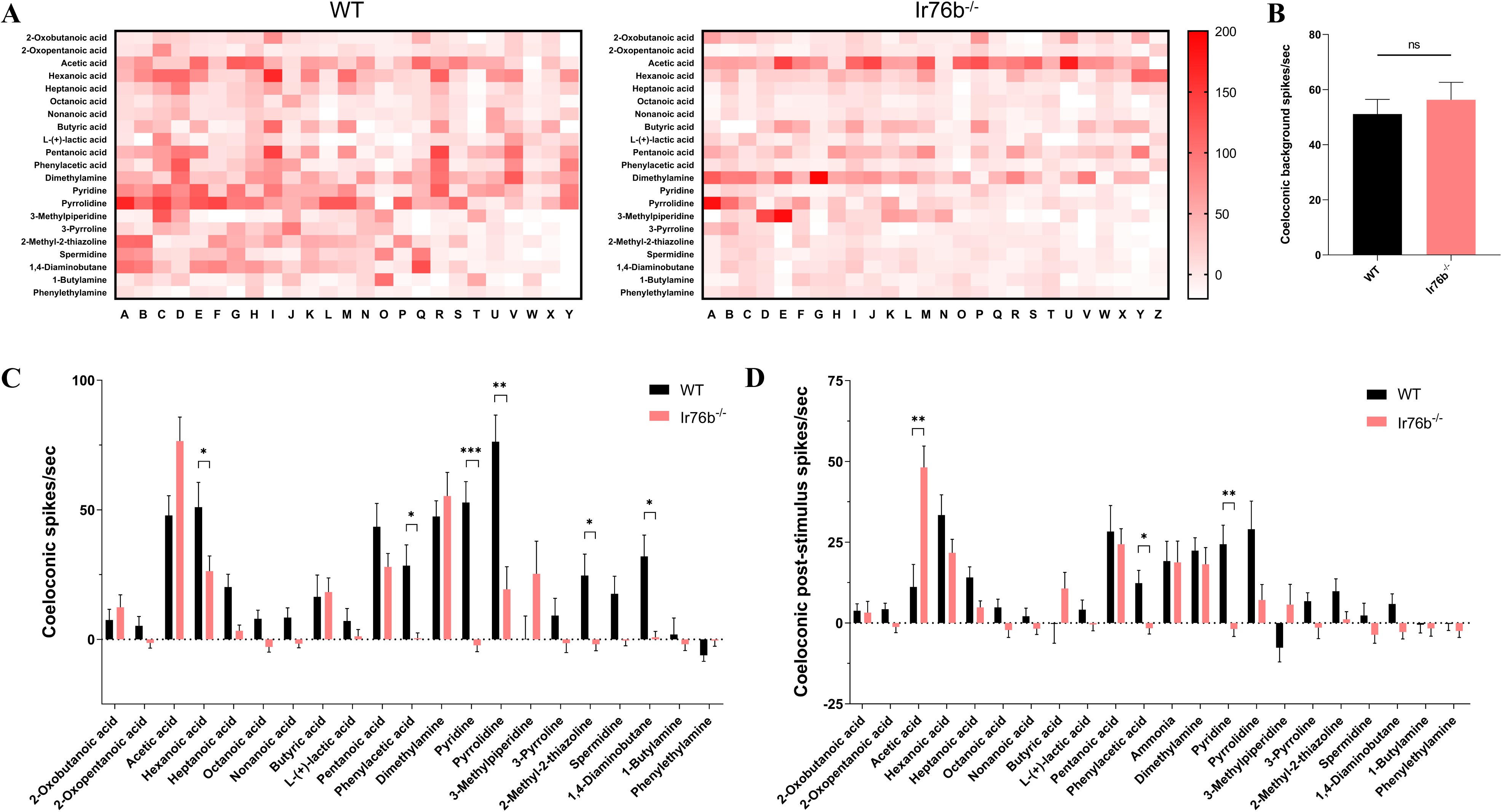
**(A)** Heatmaps showing average female coeloconic sensillum SSR responses to acids and amines in the wild-type (WT) and *AcIr76b^-/-^* (Ir76b^-/-^) females. Each column depicts a single replicate. The color scale marks the response amplitude from the highest (200 spikes/s, red) to the lowest (−20 spikes/s, white). Responses were normalized by subtracting the solvent responses. **(B)** Average female coeloconic sensillar spontaneous (background) neuronal activity. Non-parametric t-test suggests no significant differences between *AcIr76b* mutants and the wild type. **(C)** Average female coeloconic sensillum SSR responses. Multiple t-tests using Holm-Sidak method (N=25-26) suggest responses to specific acids and amines in *AcIr76b* mutants are significantly lower than in wild type. **(D)** Average female coeloconic sensillar post-stimulus “OFF” responses. Multiple t-tests using Holm-Sidak method (N=25-26) suggest post-stimulus responses to acids and amines are significantly different in *AcIr76b* mutants than in wild type. Significance levels are depicted with asterisks: p-value < 0.05 (*); p-value < 0.01 (**); p-value < 0.001 (***). Error bars = Standard error of the mean.

### Peripheral electrophysiology across accessory chemosensory appendages

To further confirm the specificity of the antennal phenotype, we carried out SSR studies on female *An. coluzzii* maxillary palp capitate pegs (cp), which are the sole sensillar class found on that appendage (5). While previous RNAseq-based transcriptome profiles of the *An. coluzzii* female maxillary palps uncovered *AcIr76b* transcripts (50, 51), no GFP-labelled cells were identified on female *Ir76b-GFP* progeny, which likely reflects the sensitivity differences between these two approaches. In that light, it was not surprising that the well-characterized GR- and OR-mediated responses of the cpA neuron to CO_2_ and the cpB/C neurons to 1-octen-3-ol and 2,4,5-trimethylthiazole, respectively (5), were not affected in *AcIr76b^-/-^* mutants (**Figure S3A**). Furthermore, in contrast to the antennae, elevated palpal responses were not observed when comparing wild-type and mutant responses using a transcuticular EPG recording of the maxillary palp (**Figure S3B**).

Lastly, in light of extensive *AcIr76b* expression across the labellum (**Figure 1G**), we also employed the ELG, an EAG/EPG-like transcuticular sampling of peripheral neuronal activity on the labellum (7), to assess responses to a panel of amines to determine whether AcIr76b might play a role in olfactory responses on that appendage. Surprisingly, in light of the robust *AcIr76b* labellum expressio these studies revealed no significant differences between the amine response profiles of the *An. coluzzii* wild-type and *AcIr76b^-/-^* female labellum (**Figure S3C**).

### *AcIr76b* expression in a distinct antennal organ in larvae

*AcIr76b* expression was also examined across the larval antennae of *An. coluzzii* which is the principal sensory appendage of this aquatic pre-adult life stage (41, 52). In addition to robust labelling of larval antennal neuronal cell bodies, which are clustered within the antennal shaft (**Figure S4A**), the *AcIr76b* promoter appears to drive GFP-labeling of dendrites that specifically innervate the sensory peg organ (**Figure S4A**). The larval sensory peg is a distinctive uniporous apical appendage on the antennae that has been hypothesized to play gustatory roles (**Figure S4B**) (52, 53). *AcIr76b* localization to the larval sensory peg dendrites is distinct from the dendritic localization of *An. coluzzii* ORs to the larval sensory cone where they are associated with olfactory signals (41, 52).

Despite their proximity at the apical tip of the larval antennae, the activity of the sensory peg is distinct from that of the OR-associated sensory cone which acts as the primary larval olfactory organ in *Anopheles* (41, 52). While technical limitations have thus far precluded direct recording of neuronal activities from the uniporous larval peg, we have nevertheless recently carried out a comprehensive electrophysiological analysis of peripheral larval sensory cone neuron responses to a wide range of volatile stimuli (41). These SSR-based methods were used to examine the functionality of wild-type and *AcIr76b^-/-^* larval sensory cones to further narrow its role to the sensory peg. As expected, based on the absence of *AcIr76b*-associated sensory cone dendrites, these recordings failed to reveal any significant differences between wild type and *AcIr76b^-/-^* mutants insofar as background neuronal activity (**Figure S4C**) or responses to a panel of amines (**Figure S4D**).

### Mating and blood-feeding deficits in *AcIr76b* mutants

In addition to their impact on the peripheral electrophysiology in adult females, *AcIr76^-/-^* mutants display a range of interesting behavioral and reproduction-related deficits and, as a result, cannot self-propagate under standard laboratory rearing protocols. To investigate this phenotype, we took advantage of a previously developed insemination-based mating bioassay (15) to reveal that self-mated *AcIr76b^-/-^* mutants display significantly impaired insemination rates when compared with their wild-type counterparts (**Figure 5A**). To investigate this phenotype further, mating studies were conducted in which *AcIr76b^-/-^* females were replaced with their wild-type counterparts. Here, wild-type females successfully mated with *AcIr76b^-/-^* males to fully rescue the mutant mating/insemination deficits. In addition, studies that paired wild-type males with *AcIr76b^-/-^* females had severe mating deficits (**Figure 5A**), indicating that *AcIr76b* plays a female-specific role in *An. coluzzii* mating.

**Figure 5.**
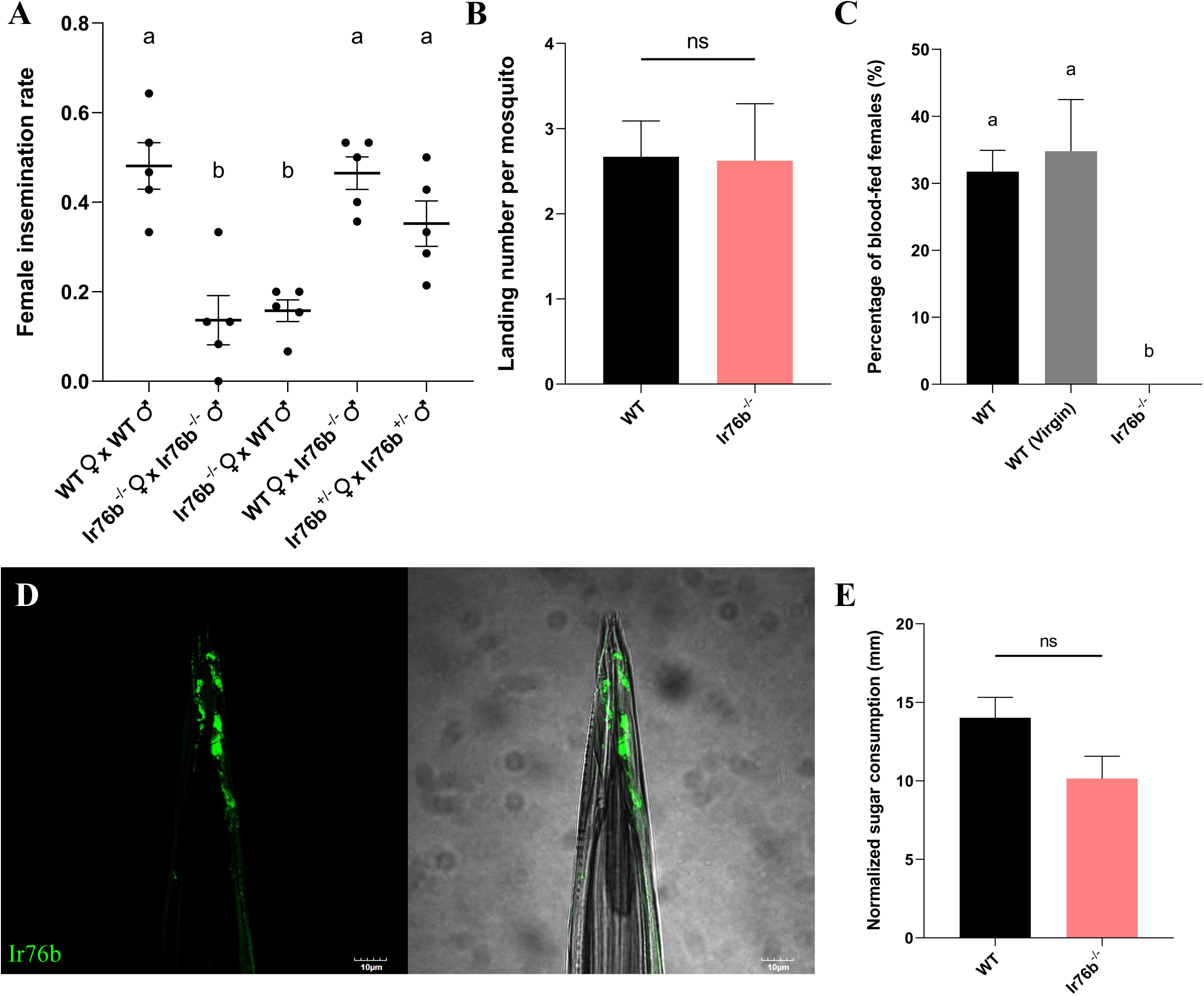
**(A)** Average insemination rate of females in different mating pairs. Mean values with different grouping letters were significantly different (N=5; one-way ANOVA; p-value < 0.05). **(B)** Average landing counts per female mosquito on the blood feeder during blood feeding. Non-parametric t-test suggests no significant differences between *AcIr76b* mutants and the wild type. **(C)** Average blood feeding rate of wild-type (WT), virgin wild-type (WT (Virgin)), and *AcIr76b^-/-^* (Ir76b^-/-^) females in the 25-min bioassay. Mean values with different grouping letters were significantly different (N=4-7; one-way ANOVA; p-value < 0.05). **(D)** A representative confocal z-stack project of female stylet showing *AcIr76b* expressions. Scale bars = 10μm. **(E)** Average sugar consumptions of female mosquitoes in the CAFE bioassay. Non-parametric t-test suggests no significant differences between *AcIr76b* mutants and the wild type. Error bars = Standard error of the mean.

We next asked what other aspects of the female reproductive pathway might underlie the mating deficits and sterility of the *AcIr76b^-/-^* mutants. To address this question, we utilized a simple digital video-based blood feeding bioassay (54) to examine whether female *AcIr76b^-/-^* mutants were able to successfully initiate and complete blood feeding. Here, 25-30 female mosquitoes were exposed to membrane feeders containing warmed blood meals supplemented with human foot odors released from worn socks and CO_2_ for 25 min. Under these conditions, approximately 30% of wild-type females successfully completed blood feeding as assessed by the presence of extended and blood-filled abdomens. We also examined alighting (landing) on the membrane feeder as a distinct component of blood-feeding behaviors by recording and quantifying those events at the surface of the blood feeder during feeding bioassays. These studies indicated that *AcIr76b^-/-^* females maintain wild-type levels of landings (**Figure 5B**), which suggests their blood-feeding deficits are not due to the reduction of host-seeking behaviors. However, under identical conditions, while wild-type levels of alighting and probing activity were observed, none (0%) of the *AcIr76b^-/-^* females that were assayed blood fed successfully (**Figure 5C**).

In light of the profound mating deficits exhibited by *AcIr76b^-/-^* females that might conceivably underlie or indirectly impact blood-feeding efficiency, we also examined the ability of virgin (unmated) wild-type females to blood feed. Here, and consistent with previous studies (55–57), wild-type virgins did not exhibit a significant alteration in blood-feeding propensity when compared with their wild-type counterparts that were first allowed to mate (**Figure 5C**).

As is the case for other mosquitoes, *Anopheles* blood feeding is a highly specialized behavior that is quite distinct from nectar (sugar) feeding (58). Subsequent to host seeking and landing, another critical aspect of mosquito blood feeding is the process of blood ingestion (uptake), which occurs through the stylet (58). The stylet is a highly specialized component of the labellum that pierces the skin of blood-meal hosts to make direct contact with blood and is used solely for blood feeding. Recent studies in *Ae. aegypti* have illustrated the roles of *Ir7a and Ir7f*, which are expressed in stylet gustatory neurons where they are specifically responsible for mediating the direct chemosensory (taste) responses to blood-meal components (59). In light of the association of the *DmIr76b* ortholog in amino acid responses in *Drosophila* (26), we hypothesized that *AcIr76b* also acts as a gustatory IR co-receptor and, in that context, would be expressed on the stylet of *An. coluzzii*. Indeed, crosses between *AcIr76b-QF2* driver and *QUAS-GFP* effector lines revealed robust expression of *AcIr76b* in the stylet (**Figure 5D**). In order to examine the blood-meal specificity of this feeding deficit phenotype, we used the well-established CAFE capillary feeding assay (15, 44) to examine sugar feeding. These studies confirmed *AcIr76b^-/-^* females can maintain wild-type levels of sugar feeding via their proboscis (**Figure 5E**) that are consistent with a highly specialized blood-feeding functionality of *AcIr76b* on the stylet of *An. coluzzii*.

## Discussion

Acting together or independently, Anopheline IRs, GRs and ORs are critical molecular components of the signal transduction processes that initiate diverse chemosensory processes underlying various elements of the mosquito lifecycle. While many of the particulars of these relationships remain opaque, it is clear the collective functionality of these elements has direct implications on the biology and the vectorial capacity of these mosquitoes. Recent advances in gene-editing approaches have led to an increasingly refined appreciation of the roles of insect IR and OR co-receptors in mediating both gustatory and olfactory signal transduction, with broad implications on mosquito behavior and physiology (21, 22, 30). Here, we have used state-of-the-art gene-editing/labelling approaches to characterize the expression and function of the ionotropic receptor co-receptor *AcIr76b* in the IR-dependent chemical ecology of the malaria vector mosquito *An. coluzzii*.

While IRs, GRs and ORs have traditionally been presumed to populate distinct classes of chemosensory neurons, recent studies demonstrated multiple types of receptors can be co-expressed in the same neuron (47, 48). Regardless of their specific cellular context, and considering the tight clustering of neurons within insect sensilla, these diverse classes of ionotropic receptors may have direct as well as indirect/modulatory roles in mediating the activation or inhibition of chemosensory neurons. For example, the *Ir25a* co-receptor is co-expressed with *Gr3* and *Orco* in *Ae. aegypti* and *Drosophila*, respectively (47, 48). In *Aedes*, knocking out *Gr3* on the CO_2_-sensitive cpA maxillary palp neuron removes CO_2_ sensitivity but importantly does not block responses to amines mediated by *AeIr25a* in the same neurons (47). In *Drosophila*, while *Orco* null mutations completely abolished OSN activity, *DmIr25a* mutations resulted in mostly non-altered or only partially reduced/increased responses suggesting that *DmIr25a* plays a regulatory role when coupled with ORs instead of acting as a canonical co-receptor (48). We initially focused on the role of *AcIr76b* in adult *An. coluzzii* females that are actively host seeking for a blood meal. In those insects, robust *Ir76b* expression occurred in the antennal grooved pegs (**Figure 1**), which are considered homologous to *Drosophila* coeloconic sensilla (60), as well as in T1 and T2 sensillum of the labellum (**Figure 1**) and across the tarsi (**Figure S1**). We also examined its expression in the larval chemosensory system which is housed on the antennae. Here, in contrast to the dendritic localization of *An. coluzzii* larval ORs, *AcIr76b* specifically labeled dendrites innervating the sensory peg, which is an apical uniporous structure associated with larval gustatory responses.

The new-found availability of CRISPR/Cas9-mediated gene targeting in *An. coluzzii* allowed us to examine loss-of-function phenotypes resulting from *AcIr76b* null mutations. These are the first reports of *Ir76b* mutagenesis in mosquitoes. In addition to defective coeloconic SSR responses to amines, consistent with that observed in *Drosophila* (24), CRISPR-generated *AcIr76b* null mutants exhibited significantly enhanced antennal responses to amines in both EAG and grooved peg SSR studies in contrast to *Ae. aegypti AeIr8a* mutants that abolish olfactory sensitivity to acids (30). These data are similar to previous studies that identified enhanced gustatory responses to sugars in *Drosophila Ir76b* mutants (61) and suggest Ir76b’s modulatory function is both context dependent and evolutionarily conserved. In *An. coluzzii*, this phenotype is restricted to the large population of non-*Orco*-expressing grooved pegs, where dramatic increases in amine responses seen in *AcIr76b* null mutants suggest that *AcIr76b* is responsible for inhibition of neuronal activity in wild-type neurons and, moreover, the presence of another IR co-receptor that is primarily responsible for action potential generation. This is likely to be Ir25a in light of its association with Ir76b orthologs in *Aedes* and *Drosophila* (18, 47). Despite the typically strong correlation between the bulk of EAG responses and grooved peg SSRs, the increased EAG responses of *AcIr76b* mutants to pyridine remain enigmatic. Taken together with the dramatic decrease in coeloconic responses of *AcIr76b* mutants to pyridine which, considering the relatively small number of coeloconic sensilla on female antennae may have a marginal impact on EAG responses, this suggests that another as-yet uncharacterized class of sensilla is responsible for this phenotype. It is intriguing to speculate that this cryptic group of sensilla may correspond to the *An. coluzzii* neurons that seem to co-express *Orco* and *Ir76b* (**Figure 1F**).

Our data provide *in vivo* evidence for the modulatory role of *AcIr76b* in antennal responses that is specific to amines and thus extend previous studies emphasizing the role of IRs in the detection and discrimination of amines (11, 62). Importantly, many compounds in this chemical class have been identified as human emanations and are components of odor blends that robustly attract blood-feeding Anopheline mosquitoes (63, 64). In contrast to exclusive gustatory functions in *Drosophila*, mosquito labella also have olfactory capabilities (7, 49). However, despite extensive *AcIr76b* expression in the labellum of female *An. coluzzii* mosquitoes, we were unable to uncover any significant electrophysiological differences to volatile odorants between wild-type and mutant mosquitoes in ELG studies. Importantly, our data do not rule out an important gustatory role for *AcIr76b* in labellum responses to contact cues such as ammonia, carboxylic and amino acids that are present in human sweat and in blood meals themselves (65, 66). Future SSR and tip-recording studies investigating the role of labellum expression of *AcIr76b* as well as similar approaches to target the gustatory responses across the labellum and tarsi of adult *An. coluzzii* which also express *AcIr76b* will doubtlessly reveal functional roles.

The restriction of larval *AcIr76b* supports a direct chemosensory role of the larval peg and, moreover, is consistent with our previous RNAi-based gene-silencing studies in *An. coluzzii* demonstrating that larval behavioral responses to aqueous butylamine are mediated by *AcIr76b* (31). While we are currently unable to physiologically examine gustatory responses of the larval sensory peg to aqueous stimuli, it is nevertheless noteworthy that *AcIr76b^-/-^* mutant larvae display wild-type responses to a panel of volatile odorants. Taken together, these data indicate that the AcIr76b-associated behavioral sensitivity of *An. Coluzzii* larvae to aqueous butylamine and perhaps other related compounds (31) is a gustatory process mediated through the sensory peg.

Over and above the expected peripheral olfactory impacts of knocking out *AcIr76b*, we have identified novel roles in Anopheline reproductive pathways. To begin, female homozygotic *AcIr76b^-/-^* mutants displayed significant reductions in insemination rates that persisted when mated with either wild-type or mutant males. In contrast, *AcIr76b^-/-^* males maintained wild-type level insemination rates when paired with wild-type females (**Figure 5A**). While these data demonstrate this phenotype is female-specific, its mechanistic basis remains unclear. We hypothesize that the Ir76b-related mating defect is linked to direct or volatile-based chemosensory processes on the labella and tarsi of *An. coluzzii* where *AcIr76b* expression is pronounced. *DmIr76b* is involved in multiple gustatory pathways in *Drosophila* (25, 26) where the labella and tarsi are recognized as gustatory appendages (44, 67, 68) and where multiple studies have identified IRs on those gustatory appendages that directly promote mating (67, 69, 70). While the chemosensory receptors involved in mosquito mating have not been identified molecularly, the tarsi of Anopheline mosquitoes are known to be essential for mating where they presumably detect contact mating pheromones (71–73). It is therefore reasonable to suggest that mating deficits observed in *AcIr76b^-/-^* females could be caused by disruption of recognition pathways that are dependent on Ir76b-mediated gustatory signaling.

In addition to the mating deficits that impede propagation of homozygous *AcIr76b^-/-^* lines, the profound absence of successful blood feeding of *AcIr76b^-/-^* females requires heterozygotic maintenance of these lines. Despite the ability of these mutants to exhibit wild-type levels of sugar feeding and, importantly, to be fully able to locate, alight on and even probe blood-containing membrane feeders that are supplemented with human foot-odor blends and CO_2_, *AcIr76b^-/-^* females fail to actually take-up (ingest) their blood meals. In *Ae. aegypti*, the labial stylets that are essential for gustatory sampling (tasting) of blood house neurons that respond to whole blood that is rich in amino acids (74) and more specifically to ATP, NaHCO_3_, and NaCl (59). More importantly, while the cognate IR co-receptors responsible for those responses remain uncharacterized, two *Aedes* IRs—*Ir7a* and *Ir7f*—acting as ‘tuning’ receptors are specifically expressed on the stylet neurons where they directly recognize those blood-specific tastant cues to activate the uptake of blood to complete blood feeding (59). In light of the robust expression of *AcIr76b* in the stylets of adult females, it is reasonable to postulate that *AcIr76b* is the gustatory IR co-receptor that is directly involved in blood tasting in *An. coluzzii*. That *AcIr76b^-/-^* mutations specifically block the uptake of blood by female mosquitoes that have located, alighted on and probed membrane feeders provides validation for this hypothesis. Inasmuch as blood feeding is paramount for *An. coluzzii* and other mosquitoes to reproduce and to acquire and transmit disease pathogens, the crucial role of *Ir76b* in this behavior makes it a provocative target for the development of novel strategies to reduce mosquito vectorial capacity.

## Author contributions

Conceived experiments: ZY, FL, HS and LJZ; Performed research: ZY, FL, HS, and AB; Analyzed data: ZY, FL, HS, and AB; Wrote the paper: ZY, FL, HS, AB, and LJZ. Approved the final manuscript: ZY, FL, HS, AB, and LJZ.

## Acknowledgements

We thank Zhen Li for mosquito rearing, Dr. H. Willi Honegger for critical comments, and all members of the Zwiebel lab for suggestions. We are very grateful to Dr. Christopher Potter (The Johns Hopkins University School of Medicine) for the generous gift of Q system mosquito lines. We thank Dr. A.M. McAinsh for scientific copy-editing and acknowledge the Vanderbilt University Cell Imaging Shared Resource Core for training and use of the Olympus FV-1000 confocal microscope. This work was conducted with the support of Vanderbilt University and a grant from the National Institutes of Health (NIAID, AI127693) to LJZ.

## References

1. M. Coetzee, et al., *Anopheles coluzzii* and *Anopheles amharicus*, new members of the *Anopheles gambiae* complex. Zootaxa 3619, 246–274 (2013).

2. A. Molina-Cruz, M. M. Zilversmit, D. E. Neafsey, D. L. Hartl, C. Barillas-Mury, Mosquito vectors and the globalization of *Plasmodium falciparum* malaria. Annu. Rev. Genet. 50, 447–465 (2016).

3. F. Van Breugel, J. Riffell, A. Fairhall, M. H. Dickinson, Mosquitoes use vision to associate odor plumes with thermal targets. Curr. Biol. 25, 2123–2129 (2015).

4. C. Montell, L. J. Zwiebel, Mosquito sensory systems. Adv. In Insect Phys. 51, 293–328 (2016).

5. T. Lu, et al., Odor coding in the maxillary palp of the malaria vector mosquito *Anopheles gambiae*. Curr. Biol. 17, 1533–1544 (2007).

6. Y. T. Qiu, J. J. A. van Loon, W. Takken, J. Meijerink, H. M. Smid, Olfactory coding in antennal neurons of the malaria mosquito, *Anopheles gambiae*. Chem. Senses 31, 845–863 (2006).

7. H. W. Kwon, T. Lu, M. Rützler, L. J. Zwiebel, Olfactory response in a gustatory organ of the malaria vector mosquito *Anopheles gambiae*. Proc. Natl. Acad. Sci. U. S. A. 103, 13526–13531 (2006).

8. F. Guidobaldi, I. J. May-Concha, P. G. Guerenstein, Morphology and physiology of the olfactory system of blood-feeding insects. J. Physiol. Paris 108, 96–111 (2014).

9. S. B. Mclver, Sensilla of mosquitoes (Diptera: Culicidae). J. Med. Entomol. 19, 489–535 (1982).

10. R. J. Pitts, A. N. Fox, L. J. Zwiebeil, A highly conserved candidate chemoreceptor expressed in both olfactory and gustatory tissues in the malaria vector *Anopheles gambiae*. Proc. Natl. Acad. Sci. U. S. A. 101, 5058–5063 (2004).

11. R. J. Pitts, S. L. Derryberry, Z. Zhang, L. J. Zwiebel, Variant ionotropic receptors in the malaria vector mosquito *Anopheles gambiae* tuned to amines and carboxylic acids. Sci. Rep. 7 (2017).

12. G. Wang, A. F. Carey, J. R. Carlson, L. J. Zwiebel, Molecular basis of odor coding in the malaria vector mosquito *Anopheles gambiae*. Proc. Natl. Acad. Sci. U. S. A. 107, 4418–4423 (2010).

13. A. F. Carey, G. Wang, C. Y. Su, L. J. Zwiebel, J. R. Carlson, Odorant reception in the malaria mosquito *Anopheles gambiae*. Nature 464, 66–71 (2010).

14. R. J. Pitts, L. J. Zwiebel, Antennal sensilla of two female Anopheline sibling species with differing host ranges. Malar. J. 5 (2006).

15. Z. Ye, et al., Ammonium transporter AcAmt mutagenesis uncovers reproductive and physiological defects without impacting olfactory responses to ammonia in the malaria vector mosquito *Anopheles coluzzii*. Insect Biochem. Mol. Biol. 134, 103578 (2021).

16. Z. Ye, et al., Heterogeneous expression of the ammonium transporter AgAmt in chemosensory appendages of the malaria vector, *Anopheles gambiae*. Insect Biochem. Mol. Biol. 120 (2020).

17. K. Sato, et al., Insect olfactory receptors are heteromeric ligand-gated ion channels. Nature 452, 1002–1006 (2008).

18. R. Benton, K. S. Vannice, C. Gomez-Diaz, L. B. Vosshall, Variant ionotropic glutamate receptors as chemosensory receptors in *Drosophila*. Cell 136, 149–162 (2009).

19. L. Abuin, et al., Functional Architecture of Olfactory Ionotropic Glutamate Receptors. Neuron 69, 44–60 (2011).

20. J. A. Butterwick, et al., Cryo-EM structure of the insect olfactory receptor Orco. Nature 560, 447–452 (2018).

21. M. Degennaro, et al., Orco mutant mosquitoes lose strong preference for humans and are not repelled by volatile DEET. Nature 498, 487–491 (2013).

22. H. Sun, F. Liu, Z. Ye, A. Baker, L. J. Zwiebel, Mutagenesis of the orco odorant receptor co-receptor impairs olfactory function in the malaria vector *Anopheles coluzzii*. Insect Biochem. Mol. Biol. 127 (2020).

23. C. A. Yao, R. Ignell, J. R. Carlson, Chemosensory coding by neurons in the coeloconic sensilla of the *Drosophila* antenna. J. Neurosci. 25, 8359–8367 (2005).

24. A. Vulpe, P. Mohapatra, K. Menuz, Functional characterization of odor responses and gene expression changes in olfactory co-receptor mutants in *Drosophila*. bioRxiv (2021) https://doi.org/10.1101/2021.06.18.449017 (June 26, 2021).

25. Y. V. Zhang, J. Ni, C. Montell, The molecular basis for attractive salt-taste coding in *Drosophila*. Science 340, 1334–1338 (2013).

26. A. Ganguly, et al., A molecular and cellular context-dependent role for Ir76b in detection of amino acid taste. Cell Rep. 18, 737–750 (2017).

27. L. Ni, et al., The ionotropic receptors IR21a and IR25a mediate cool sensing in *Drosophila*. Elife 5 (2016).

28. A. Enjin, et al., Humidity sensing in *Drosophila*. Curr. Biol. 26, 1352–1358 (2016).

29. S. Min, M. Ai, S. A. Shin, G. S. B. Suh, Dedicated olfactory neurons mediating attraction behavior to ammonia and amines in *Drosophila*. Proc. Natl. Acad. Sci. U. S. A. 110, 1321–1329 (2013).

30. J. I. Raji, et al., *Aedes aegypti* mosquitoes detect acidic volatiles found in human odor using the IR8a pathway. Curr. Biol. 29, 1253–1262.e7 (2019).

31. C. Liu, et al., Distinct olfactory signaling mechanisms in the malaria vector mosquito *Anopheles gambiae*. PLoS Biol. 8, 27–28 (2010).

32. C. J. Potter, B. Tasic, E. V. Russler, L. Liang, L. Luo, The Q system: A repressible binary system for transgene expression, lineage tracing, and mosaic analysis. Cell 141, 536–548 (2010).

33. A. N. Fox, R. J. Pitts, H. M. Robertson, J. R. Carlson, L. J. Zwiebel, Candidate odorant receptors from the malaria vector mosquito *Anopheles gambiae* and evidence of down-regulation in response to blood feeding. Proc. Natl. Acad. Sci. U. S. A. 98, 14693–14697 (2001).

34. E. Suh, D. H. Choe, A. M. Saveer, L. J. Zwiebel, Suboptimal larval habitats modulate oviposition of the malaria vector mosquito *Anopheles coluzzii*. PLoS One 11 (2016).

35. F. Liu, Z. Ye, A. Baker, H. Sun, L. J. Zwiebel, Gene editing reveals obligate and modulatory components of the CO_2_ receptor complex in the malaria vector mosquito, *Anopheles coluzzii*. Insect Biochem. Mol. Biol. 127 (2020).

36. A. Hammond, et al., A CRISPR-Cas9 gene drive system targeting female reproduction in the malaria mosquito vector *Anopheles gambiae*. Nat. Biotechnol. 34, 78–83 (2016).

37. K. Labun, et al., CHOPCHOP v3: Expanding the CRISPR web toolbox beyond genome editing. Nucleic Acids Res. 47, W171–W174 (2019).

38. E. Pondeville, et al., Efficient φc31 integrase-mediated site-specific germline transformation of *Anopheles gambiae*. Nat. Protoc. 9, 1698–1712 (2014).

39. B. J. Matthews, M. A. Younger, L. B. Vosshall, The ion channel ppk301 controls freshwater egg-laying in the mosquito *Aedes aegypti*. Elife 8 (2019).

40. F. Liu, L. Chen, A. G. Appel, N. Liu, Olfactory responses of the antennal trichoid sensilla to chemical repellents in the mosquito, *Culex quinquefasciatus*. J. Insect Physiol. 59, 1169–1177 (2013).

41. H. Sun, F. Liu, A. Baker, L. J. Zwiebel, Neuronal odor coding in the larval sensory cone of *Anopheles coluzzii*: Complex responses from a simple system. bioRxiv (2020) https://doi.org/10.1101/2020.09.09.290544.

42. C. J. Den Otter, M. Behan, F. W. Maes, Single cell responses in female *Pieris brassicae* (Lepidoptera: Pieridae) to plant volatiles and conspecific egg odours. J. Insect Physiol. 26, 465–472 (1980).

43. R. J. Pitts, C. Liu, X. Zhou, J. C. Malpartida, L. J. Zwiebel, Odorant receptor-mediated sperm activation in disease vector mosquitoes. Proc. Natl. Acad. Sci. U. S. A. 111, 2566–2571 (2014).

44. E. J. Dennis, O. V. Goldman, L. B. Vosshall, *Aedes aegypti* mosquitoes use their legs to sense DEET on contact. Curr. Biol. 29, 1551–1556.e5 (2019).

45. O. Riabinina, et al., Organization of olfactory centres in the malaria mosquito *Anopheles gambiae*. Nat. Commun. 7 (2016).

46. F. Diao, B. H. White, A novel approach for directing transgene expression in *Drosophila*: T2A-Gal4 in-frame fusion. Genetics 190, 1139–1144 (2012).

47. M. A. Younger, et al., Non-canonical odor coding ensures unbreakable mosquito attraction to humans. bioRxiv (2020) https://doi.org/10.1101/2020.11.07.368720.

48. D. Task, et al., Widespread polymodal chemosensory receptor expression in *Drosophila* olfactory neurons. bioRxiv (2020) https://doi.org/10.1101/2020.11.07.355651.

49. A. M. Saveer, R. J. Pitts, S. T. Ferguson, L. J. Zwiebel, Characterization of chemosensory responses on the labellum of the malaria vector mosquito, *Anopheles coluzzii*. Sci. Rep. 8 (2018).

50. R. J. Pitts, D. C. Rinker, P. L. Jones, A. Rokas, L. J. Zwiebel, Transcriptome profiling of chemosensory appendages in the malaria vector *Anopheles gambiae* reveals tissue- and sex-specific signatures of odor coding. BMC Genomics 12 (2011).

51. G. Athrey, et al., Chemosensory gene expression in olfactory organs of the anthropophilic *Anopheles coluzzii* and zoophilic *Anopheles quadriannulatus*. BMC Genomics 18 (2017).

52. Y. Xia, et al., The molecular and cellular basis of olfactory-driven behavior in *Anopheles gambiae* larvae. Proc. Natl. Acad. Sci. U. S. A. 105, 6433–6438 (2008).

53. D. Nicastro, R. R. Melzer, H. Hruschka, U. Smola, Evolution of small sense organs: Sensilla on the larval antennae traced back to the origin of the Diptera. Naturwissenschaften 85, 501–505 (1998).

54. C. J. McMeniman, R. A. Corfas, B. J. Matthews, S. A. Ritchie, L. B. Vosshall, Multimodal integration of carbon dioxide and other sensory cues drives mosquito attraction to humans. Cell 156, 1060–1071 (2014).

55. J. D. Charlwood, et al., “A mate or a meal” - pre-gravid behaviour of female *Anopheles gambiae* from the islands of São Tomé and Príncipe, West Africa. Malar. J. 2, 1–11 (2003).

56. T. W. Scott, W. Takken, Feeding strategies of anthropophilic mosquitoes result in increased risk of pathogen transmission. Trends Parasitol. 28, 114–121 (2012).

57. C. M. Stone, I. M. Hamilton, W. A. Foster, A survival and reproduction trade-off is resolved in accordance with resource availability by virgin female mosquitoes. Anim. Behav. 81, 765–774 (2011).

58. R. M. Gordon, W. H. R. Lumsden, A study of the behaviour of the mouth-parts of mosquitoes when taking up blood from living tissue; together with some observations on the ingestion of microfilariae. Ann. Trop. Med. Parasitol. 33, 259–278 (1939).

59. V. Jové, et al., Sensory discrimination of blood and floral nectar by *Aedes aegypti* mosquitoes. Neuron 108, 1163–1180.e12 (2020).

60. A. Ray, Reception of odors and repellents in mosquitoes. Curr. Opin. Neurobiol. 34, 158–164 (2015).

61. H. L. Chen, U. Stern, C. H. Yang, Molecular control limiting sensitivity of sweet taste neurons in *Drosophila*. Proc. Natl. Acad. Sci. U. S. A. (2019) https://doi.org/10.1073/pnas.1911583116.

62. A. Hussain, et al., Ionotropic chemosensory receptors mediate the taste and smell of polyamines. PLoS Biol. 14 (2016).

63. C. K. Mweresa, et al., Enhancing attraction of African malaria vectors to a synthetic odor blend. J. Chem. Ecol. 42, 508–516 (2016).

64. R. I. Ellin, et al., An apparatus for the detected and quantitation of volatile human effluents. J. Chromatogr. A 100, 137–152 (1974).

65. L. B. Baker, Physiology of sweat gland function: The roles of sweating and sweat composition in human health. Temperature 6, 211–259 (2019).

66. R. C. Burke, T. H. Lee, V. Buettner-Janusch, Free amino acids and water soluble peptides in stratum corneum and skin surface film in human beings. Yale J. Biol. Med. 38, 355–373 (1966).

67. C. Montell, A taste of the *Drosophila* gustatory receptors. Curr. Opin. Neurobiol. 19, 345–353 (2009).

68. J. T. Sparks, B. T. Vinyard, J. C. Dickens, Gustatory receptor expression in the labella and tarsi of *Aedes aegypti*. Insect Biochem. Mol. Biol. 43, 1161–1171 (2013).

69. Z. He, Y. Luo, X. Shang, J. S. Sun, J. R. Carlson, Chemosensory sensilla of the *Drosophila* wing express a candidate ionotropic pheromone receptor. PLoS Biol. (2019) https://doi.org/10.1371/journal.pbio.2006619.

70. T. W. Koh, et al., The *Drosophila* IR20a clade of ionotropic receptors are candidate taste and pheromone receptors. Neuron 83, 850–865 (2014).

71. A. Diabate, F. Tripet, Targeting male mosquito mating behaviour for malaria control. Parasites and Vectors 8 (2015).

72. L. S. Baik, J. R. Carlson, The mosquito taste system and disease control. Proc. Natl. Acad. Sci. U. S. A. 117, 32848–32856 (2020).

73. J. D. Charlwood, M. D. R. Jones, Mating behaviour in the mosquito, Anopheles gambiae s.l. I. Close range and contact behaviour. Physiol. Entomol. 4, 111–120 (1979).

74. A. M. Proenza, A. Palou, P. Roca, Amino acid distribution in human blood: A significant pool of amino acids is adsorbed onto blood cell membranes. Biochem. Mol. Biol. Int. 34, 971–982 (1994).

