## Supplemental Information for "Discrete Roles of the Ir76b Ionotropic Co-Receptor Impact Olfaction, Blood Feeding, and Mating in the Malaria Vector Mosquito *Anopheles coluzzii*"

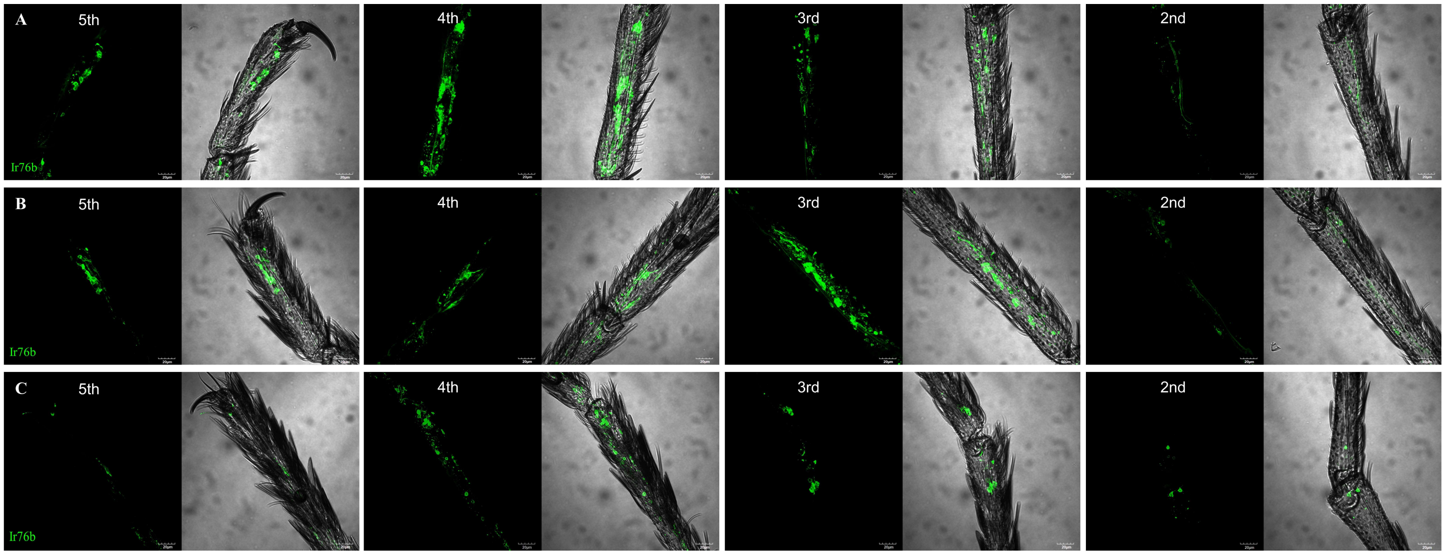


**Figure S1.** Representative confocal z-stack projects of female tarsi showing *AcIr76b* expressions in 2nd-5th segments of **(A)** pro-, **(B)** meso-, and **(C)** meta-tarsi. Scale bars = 20μm.

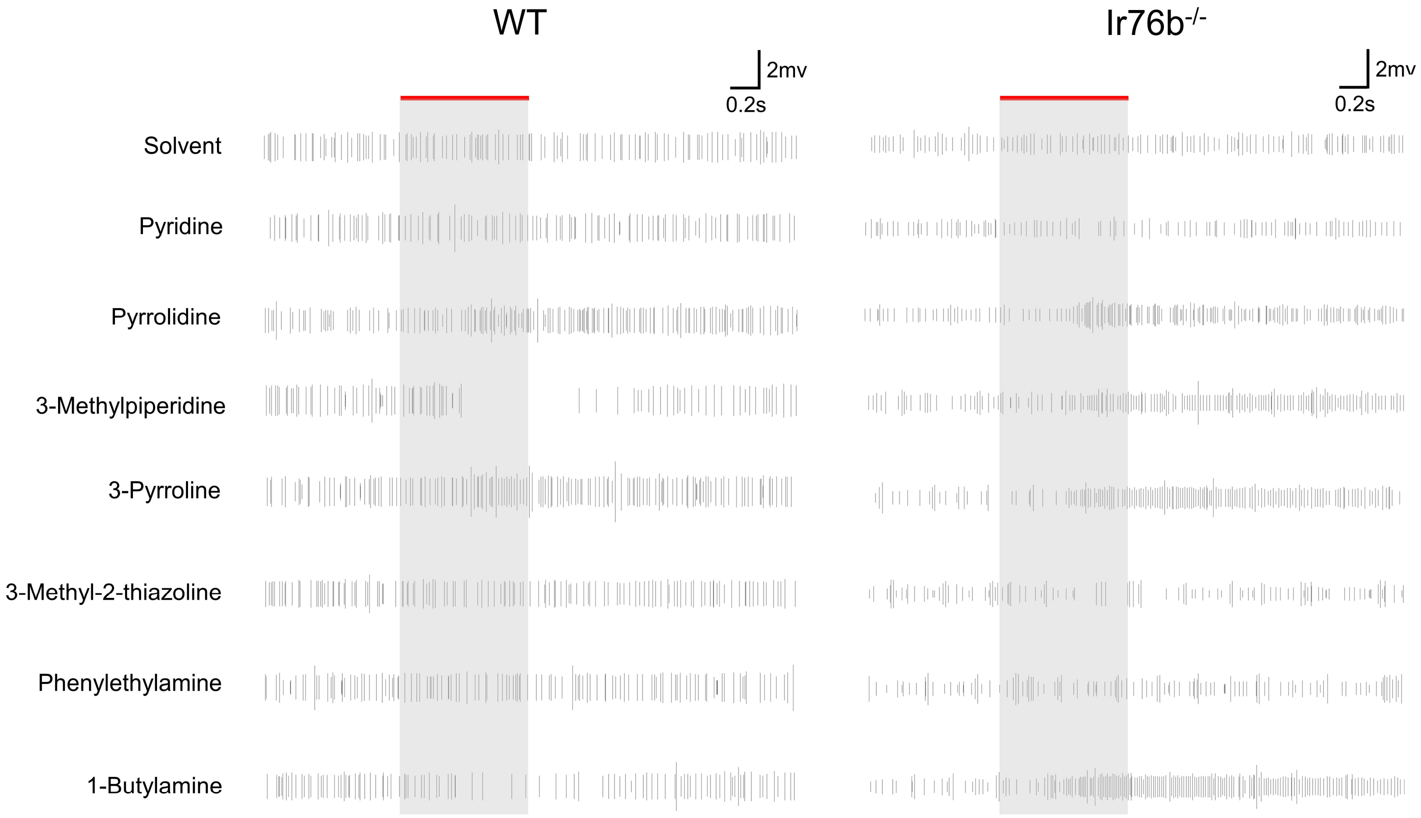


**Figure S2.** Representative grooved peg SSR signals to amines in the wild-type (WT) and *AcIr76b*$^{-/-}$ (Ir76b$^{-/-}$) females. Action potential waveforms are simplified into spikes. The red bars indicate duration of stimulations (0.5s).

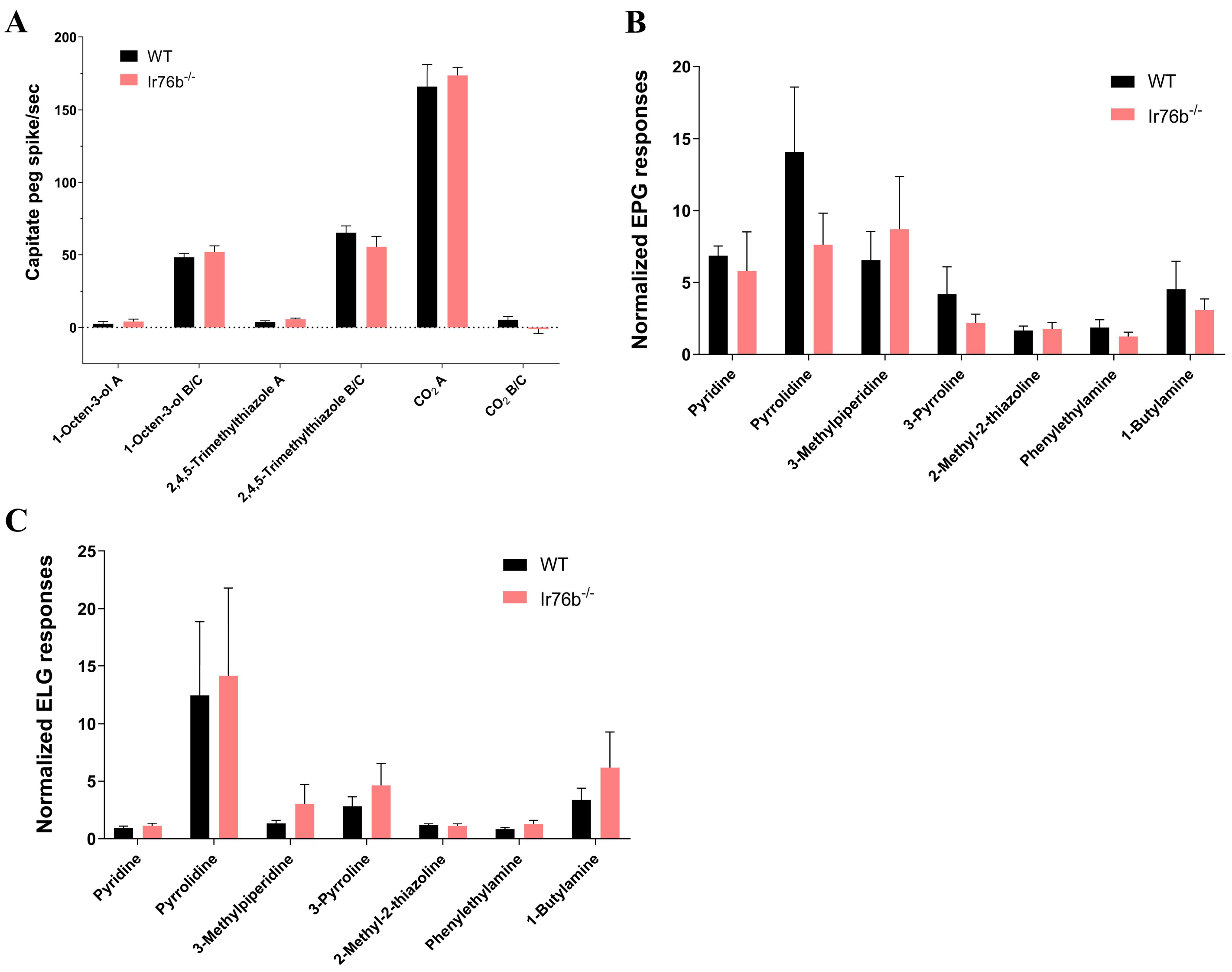


**Figure S3. (A)** Bar plot showing the means of female capitate peg SSR responses in cpA (A) and cpB/C (B/C) neurons. Multiple t-tests using Holm-Sidak method (N=5) suggest no significant differences between *AcIr76b* mutants and the wild type. **(B)** Average EPG responses to amines. Multiple t-tests using Holm-Sidak method (N=6) suggest no significant differences between *AcIr76b* mutants and the wild type. Responses were normalized to the solvent responses. **(C)** Average ELG responses to amines. Multiple t-tests using Holm-Sidak method (N=6-7) suggest no significant differences between *AcIr76b* mutants and the wild type. Responses were normalized to the solvent responses. Error bars = Standard error of the mean.

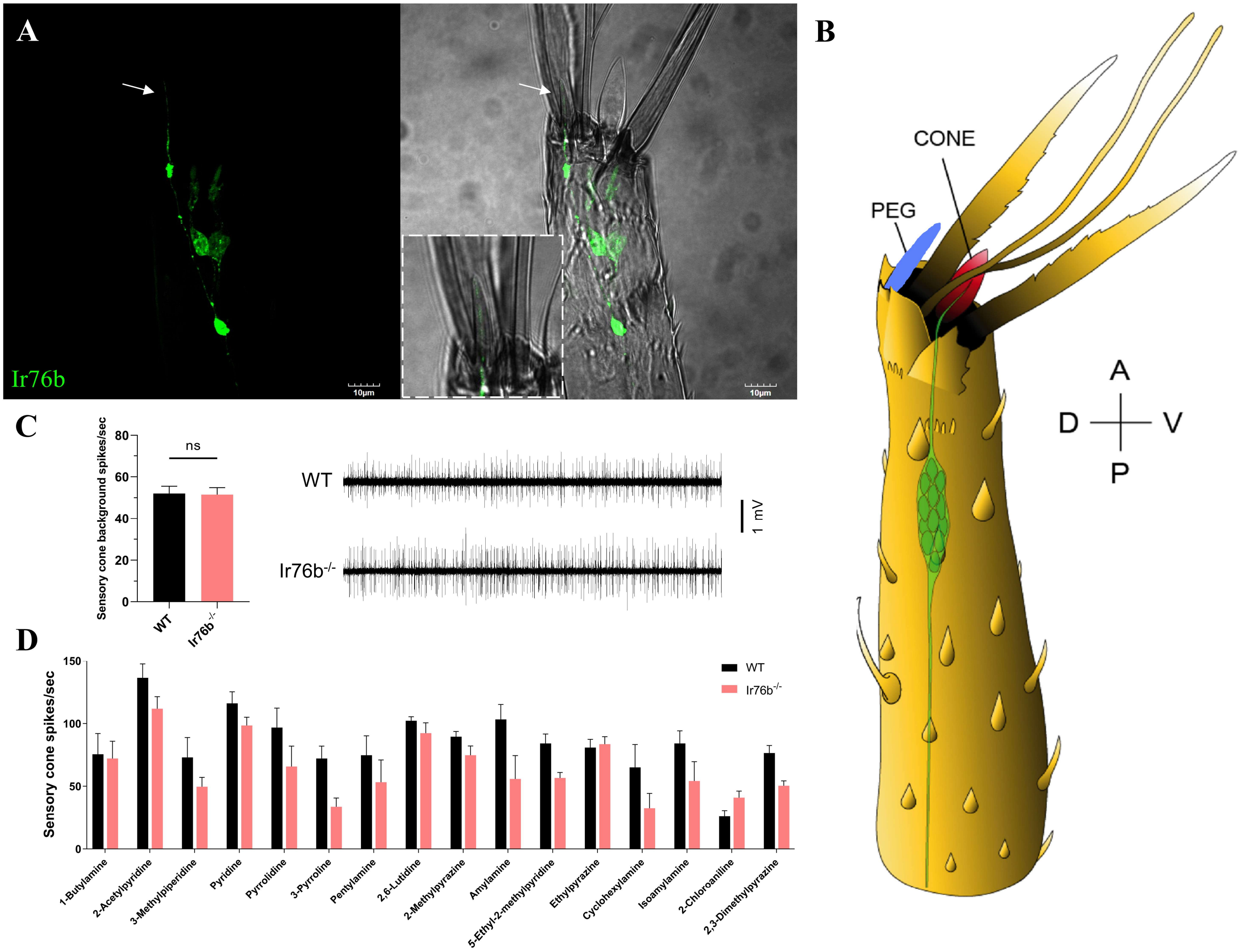


**Figure S4. (A)** A representative confocal z-stack project of a larval antenna showing *AcIr76b* is expressed in dendrites innervating the peg (highlighted by an arrow and enlarged within dashed lines). Scale bars = 10µm. **(B)** Schematics of *An. coluzzii* larval antennal structure highlighting the peg. **(C)** Average spontaneous (background) activity and representative SSR traces from larval sensory cone. Non-parametric t-tests reveal no significant differences between *AcIr76b* mutants and the wild type. **(D)** Average SSR responses in larval sensory cone to a panel of amines. Multiple t-tests using Holm-Sidak method (N=6-11) reveal no significant differences between *AcIr76b* mutants and the wild type. Error bars = Standard error of the mean.

**Table S1.** List of primers (5'→3') used in mosquito line generations and validations.

| | |
|---|---|
| Mutagenesis | |
| Ir76b_gRNA_F | TGCTGTCCACACGTCCCGATTGAAG |
| Ir76b_gRNA_R | AAACCTTCAATCGGGACGTGTGGAC |
| Ir76b_AarI_F | GGTACACCTGCGCAGTCGCGTTAAGCCGTGCGCAAGAAG |
| Ir76b_AarI_R | GGACCACCTGCCCTCTTATAAGGGGGCGAGCAGACCGGA |
| Ir76b_SapI_F | GAGTGCTCTTCTTATCAATCGGGACGTGTGGATAC |
| Ir76b_SapI_R | GGCTGCTCTTCGGACAATTATGTATGGCGACGACGA |
| T2A-QF | |
| Ir76b_LArm_F | TTCGAGCTCGGTACCCGGGGATCCTTGCCGAGCGATGGCCCGA |
| Ir76b_LArm_R | TCTGCCCTCTCCGGGGGCGAGCAGACCGGA |
| Ir76b_T2A_F | TCTGCTCGCCCCCGGAGAGGGCAGAGGAAGTC |
| Ir76b_T2A_R | CACGTCCCGATTGTAAGATACATTGATGAGTTTGGACAAAC |
| Ir76b_RArm_F | CAATGTATCTTACAATCGGGACGTGTGGATAC |
| Ir76b_RArm_R | AAGCTTGCATGCCTGCAGGTCGACTAACTTGAGCGTGCTCACC |
| PCR validation | |
| Ir76b_F | ATGGAAAAGTTCAACTTCACC |
| Ir76b_R | CTCCCGTTGGTGACAAGGT |
